# Generating fast-twitch myotubes *in vitro* using an optogenetic-based, quantitative contractility assay

**DOI:** 10.1101/2022.10.04.510824

**Authors:** Katharina Hennig, David Hardman, David Barata, Inês Martins, Miguel O. Bernabeu, Edgar R. Gomes, William Roman

## Abstract

The composition of fiber types within skeletal muscle impacts the tissue’s physiological characteristics and susceptibility to disease and ageing. *In vitro* systems should therefore account for fiber type composition when modelling muscle conditions. To induce fiber specification *in vitro*, we designed a quantitative contractility assay based on optogenetics and particle image velocimetry. We submitted cultured myotubes to long-term intermittent light stimulation patterns and characterized their structural and functional adaptations. After several days of *in vitro* exercise, myotubes contract faster and are more resistant to fatigue. The enhanced contractile functionality was accompanied by advanced maturation such as increased width and upregulation of neuron receptor genes. We observed an upregulation in the expression of distinct myosin heavy chain isoforms (namely, neonatal-Myh8 and fast-Myh), which induced a shift towards a fast fiber phenotype. This long-term *in vitro* exercise strategy can be used to study fiber specification and refine muscle disease modelling.

## Introduction

Skeletal muscle tissue possesses distinct muscle fiber types, which are broadly categorized by their contractile properties: slow (type I) and fast (type II) twitch muscle cells (myofibers). The proportion of fiber types depends on the functional demand of specific muscles (Agbulut et al., 2003). Slow, fatigue-resistant fibers are predominant in muscles involved in posture whereas fast twitching fibers possess a higher mechanical force output and are present in muscles devoted to motion (Schiaffino et al., 2011). Moreover, muscle composition can adapt through fiber type switching in response to internal and external factors, such as neuromuscular activity, exercise, or hormone exposure (Blaauw et al., 2013). Ageing, injury and disease have also been linked to changes in fiber type composition (Jansen & Fladby, 1990). For example, sarcopenia, the age-related loss of muscle mass, is characterized by a loss of type II-associated satellite cells (Frontera & Ochala, 2015) whereas nerve denervation, which occurs in certain neuromuscular disorders, leads to a slow-to-fast fiber switch (Peggion et al., 2017; Ciciliot et al., 2013). Muscle disorders also tend to predominantly affect one muscle type over another such as in Duchenne Muscular Dystrophy (DMD) in which type II myofibers are more susceptible to damage (Talbot & Maves, 2016).

Despite the importance of fiber type in disease pathophysiology, fiber type is rarely considered when modelling muscle disorders *in vitro* (Guo et al., 2012). Advances in tissue engineering and *in vitro* cell models have offered new tools to better mimic native muscle tissue and muscle disorders. Various techniques have contributed to enhance the structural maturity and contractile functionality of artificial muscle such as: (i) cell alignment via microfabrication (Bajaj et al., 2011; Zhao et al., 2009), (ii) 3D culture systems (Maffioletti et al., 2018), (iii) exercise-like stimuli (Aguilar-Agon et al., 2019; Sebille et al., 2017; Asano et al., 2015), and (iv) integration of other cell types (e.g. innervating neurons (Osaki et al., 2018; Guo et al., 2020; Bakooshli et al., 2019) and vasculature (Osaki et al., 2018)). The validation of improved myogenesis relies on a combination of molecular, morphological, and functional assays. However, most studies do not perform any characterization of fiber type composition. As these *in vitro* systems are increasingly adopted for drug screening and biological investigation, the systematic characterization of muscle tissue composition is crucial, especially to model diseases affecting distinct fiber types.

The expression of distinct myosin isoforms determines fiber type and contractile properties. Myosin consists of 2 myosin heavy chains (Myh), 2 regulatory light chains and 2 essential chains. Myh with its ATPase activity and actin-binding domain is responsible for power strokes during contractions. Interestingly, ATPase activity of distinct Myh isoforms is directly linked with the myofiber’s contraction velocity (Bottinelli et al., 1994; Bottinelli et al., 1991). Hence, Myh isoforms are the most appropriate biomarkers to characterize fiber types. During development, the expression of Myh isoforms follows a sequential, temporal pattern (Agbulut et al., 2003): developmental Myh, namely embryonic Myh3 (emb-Myh3) and neonatal Myh8 (neo-Myh8), are expressed together with slow Myh7 (slow-Myh7). Throughout the course of differentiation, developmental Myh isoforms disappear when slow-Myh7 or fast Myh isoform (fast-Myh1 and fast-Myh2) expression is upregulated. The predominantly expressed Myh isoform determines the adult fiber type and its associated contraction dynamics. Compared to adult Myh isoforms, embryonic Myh isoforms possess slower activation and relaxation kinetics and a lower rate of force production (Racca et al., 2013), while adult fibers that predominately express slow-Myh7 contract with a slower velocity than fibers containing fast Myh isoforms (Pellegrino et al., 2013; Johnson et al., 2019).

In this study, we describe an experimental strategy to alter fiber type composition of *in vitro* muscle systems. By subjecting myotubes to long-term, intermittent mechanical training, we monitored structural, functional, and molecular changes underlying fiber type switching. Long-term *in vitro* exercise alters the temporal protein expression pattern of developmental and adult Myh isoforms: trained myotubes upregulate the expression of neo-Myh8 and fMyh at day 4 of differentiation. Our results suggest that rhythmic, long-term mechanical training triggers a phenotypical shift toward a fatigue-resistant, fast fiber type. This approach could be used to model fiber type-specific diseases and identify new therapeutic targets.

## Results

### Designing a quantitative, optogenetics-based *in vitro* contractility assay

We aimed to determine if long-term exercise of *in vitro* myotubes alters contraction dynamics. To do so, we relied on an *in vitro* muscle protocol, which generates highly differentiated myotubes (Falcone et al., 2014; Pimentel et al., 2017). After 7 days of differentiation, these myotubes exhibit hallmarks of mature muscle cells, such as peripheral nuclei, striations, and contractility. To control their contraction, we infected these cultures with the AAV9-pACAGW-ChR2-Venus (adenovirus) at day 0, resulting in a high percentage of myotubes (90.38 %; **supplementary figure 1A, B**) expressing ChR2 at day 4. We confirmed the cells’ functional photosensitivity by exposing ChR2-expressing myotubes to blue light under an inverted fluorescent microscope. This resulted in 86.99 % of total myotubes performing cycles of contraction under continuous illumination.

To submit myotubes to specific and long-term stimulation patterns, we designed an optogenetic-based, *in vitro* training platform (termed OptoPlate; **figure 1A, B**). The OptoPlate emits blue light pulses with high temporal control for extended durations and is compatible with cell culture. The device consists of a 3D printed 6-well plate made of polylactic acid (PLA) with integrated blue LEDs (475 nm). Seven LEDs are grouped under each well and can be controlled independently via an Arduino board. A touch screen interface allows to set training parameters (e.g. pulse length, light intensity, and duration) for each 35 mm dish separately.

**Figure 1).**
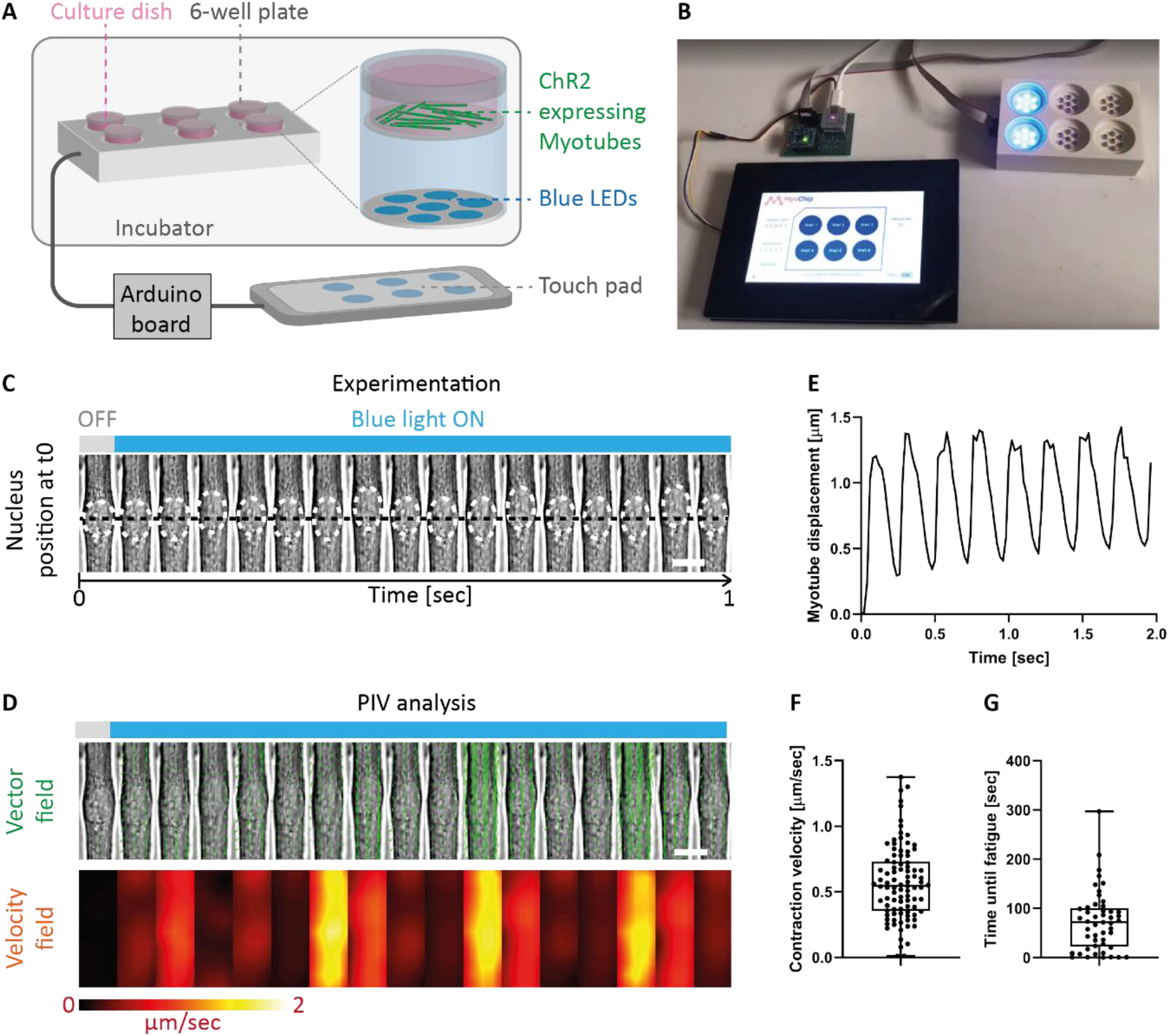
Strategy to submit myotubes to long-term light-induced exercise and quantify contraction kinetics with single cell resolution. **A, B)** Schematic illustration and image of designed OptoPlate. **C)** Kymograph of a 4-day myotube continuously stimulated with blue light. White dashed circles highlight the nucleus and the black dashed line marks the initial nuclear position before stimulation. **D)** Particle image velocimetry (PIV) analysis applied to myotube displacement over time. Displacement vectors (green arrows) and velocity fields (heat map) convey myotube movement. **F)** Graph plotting computed displacement curve for a single contracting myotube. **G)** Boxplot of myotube contraction velocity averaged over 2 seconds. **H)** Boxplot of time needed for cells to enter fatigue when continuously light stimulated. Scale bars: 10 μm.

To measure contraction dynamics after long-term training, we designed a quantitative, functional contractility assay based on particle image velocimetry (PIV) analysis. We experimentally assessed contractile behaviors at day 4 of differentiation (after 3 days of training) by continuously illuminating ChR2-expressing myotubes under the microscope and recording contractions for 2 seconds (**figure 1C**). We used PIVlab, an imaged-based PIV analysis software (Thielicke et al., 2014), to analyze contraction kinetics quantitatively. The program divides images into interrogation windows, computes the local displacement of two consecutive windows via maximum correlation methods and outputs vector fields and velocity magnitudes (**figure 1D**). We use both these measurements to determine contraction kinetics such as myotube displacement and contraction velocities of single myotubes (**figure 1E, F**). Finally, we monitored the time until myotubes enter fatigue by continuously stimulating cultures with blue light until they stop responding (**figure 1G**). We employed this optogenetic contractility assay to evaluate the effect of long-term exercise on muscle cell function.

### Long-term *in vitro* exercise accelerates myotube contraction speed and decreases fatigability

The OptoPlate allows to submit myotubes to distinct long-term exercise programs by setting specific light stimulation parameters (intensity, duration, and frequency). To trigger contractions in a non-invasive manner with high temporal control, myotubes were infected with AAV9-pACAGW-ChR2-Venus at day 0 of differentiation (**figure 2A**). We first tested if an optimal stimulation frequency enhances contractile properties of myotubes. Three OptoTraining protocols were tested: 50 ms light pulses at 2, 5 or 10 Hz (**figure 2B**). To mimic chronic exercise, we trained myotubes for 8 hours per day.

**Figure 2).**
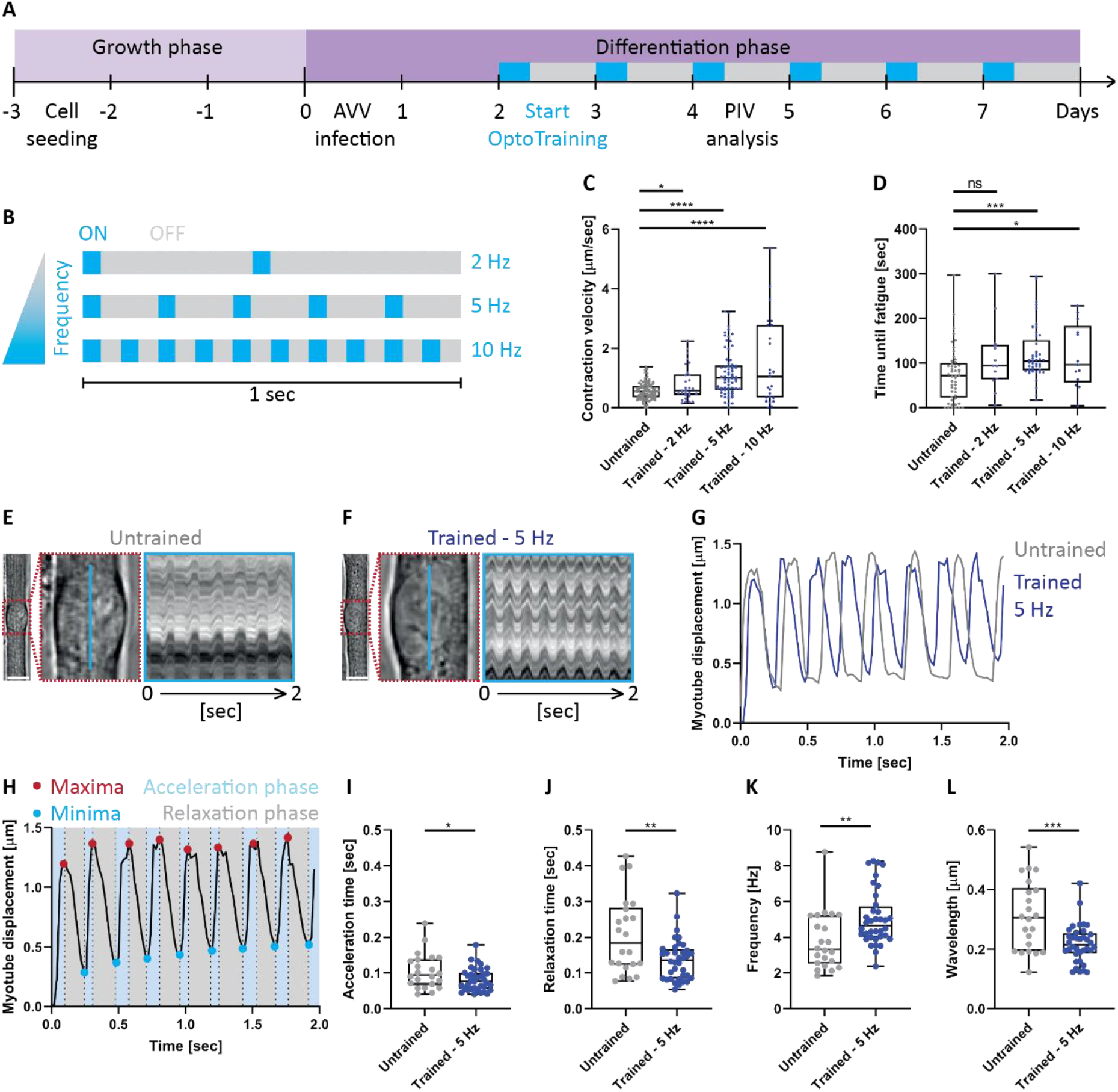
Trained myotubes undergo faster contraction cycles. **A**) Experimental protocol to submit in vitro primary mouse myotubes to OptoTraining. After the initial growth phase, myoblasts differentiate into multinucleated myotubes. Myotubes were infected with AAV9-pACAGW-ChR2-Venus at day 0 and submitted to OptoTraining from day 2 (8 hours per day, blue boxes). **B)** Schematic showing the three OptoTraining protocols at 2, 5 or 10 Hz. **C, D)** Boxplots depicting contraction velocities **(C)** and time until fatigue **(D)** of myotubes after 3 days of OptoTraining with different frequencies. **E, F)** Left: Representative brightfield images of single myotubes with magnifications of myonuclei (red dashed boxes). Right: 2 second kymographs (blue line) of untrained **(E)** and trained **(F)** myotubes when continuously stimulated with blue light. **G)** Single track displacement curves of untrained (grey) and trained (5 Hz light stimulation frequency: blue) myotubes. **H)** Representative displacement curve illustrating metrics of quantitative analysis during contraction: detection of maxima (red) and minima (blue) allows to dissect the curve into acceleration (light blue) and relaxation (grey) phases. **I, J)** Acceleration and relaxation time of individual myotubes averaged over contractions over 2 seconds. **K, L)** Frequency and wavelength for untrained and trained myotubes. Scale bars: 10 μm.

By using low light intensity, we minimized any phototoxicity effect. Additionally, long-term OptoTraining did not cause any sarcomeric damage. Although both untrained and trained myotubes exhibit local sarcomeric scars visualized via filamin C staining (Roman et al., 2021), they display no significant increase in filamin C mRNA or protein expression (**supplementary figure 2A-E**).

Using our PIV-based contractility assay, we investigated functional adaptations of myotubes to the different Optotraining protocols. Interestingly, myotube contraction velocity positively correlates with light stimulation frequency: higher stimulating frequencies resulting in faster myotube twitching after training (**figure 2C**). However, 10 Hz stimulated myotubes are more prone to fatigue (**figure 2D**) for a marginal increase in contraction velocity when compared to the 5 Hz cohort. As such, we selected 5 Hz as the optimal training frequency. All further experiments compare untrained control myotubes to the 5 Hz OptoTraining protocol (from here on generally referred to as “trained myotubes”).

Individual trained myotubes possess faster contraction cycles compared to untrained control myotubes (**figure 2E-G**). To understand the origin of this increased contraction speed, we performed a deeper quantitative analysis of contraction kinetics. For this, we developed an algorithm dissecting individual displacement curves of contracting myotubes into acceleration and relaxation phases (**figure 2H**) and observed a significantly shorter acceleration (**figure 2I**) and relaxation phase for trained myotubes (**figure 2J**). These trained myotubes therefore undergo contractions with higher frequency and shorter wavelength leading to faster contraction cycles (**figure 2K, L**).

### OptoTraining increases cell width and promotes synaptogenesis

Since contraction depends on muscle architecture, we sought to understand if changes in cellular structures underlie the faster contraction cycles observed after long-term *in vitro* training. We performed a quantitative structural analysis by extracting parameters of myotube maturation from immunofluorescent images at day 4 of differentiation (**figure 3A, B**; Hardman et al., 2021). Trained myotubes display an increased cell width (**figure 3C**), suggesting an addition of sarcomeres due to training. We also observed shorter myonuclear distances (**figure 3D**) whereas the myonuclei variability coefficient (a measure of how uniformly myonuclei are distributed within myotubes) remains unchanged (**figure 3E**). These maturation trends are conserved for 2 and 10 Hz stimulation frequencies albeit more discrete (**supplementary figure 3A-C**). Surprisingly, we did not observe a significant difference in the percentage of striated myotubes at day 4 (**figure 3F**), suggesting that initiation of myofibril assembly is similar in trained and untrained cultures. However, western blot analysis shows an increased protein expression of α-actinin for trained myotubes which persists over 7 days (**figure 3G-I**). Nevertheless, we did not find a correlation between fiber width and contraction speed **(figure 3J)** indicating that faster contractile dynamics in trained cultures are not due to thicker myotubes.

**Figure 3).**
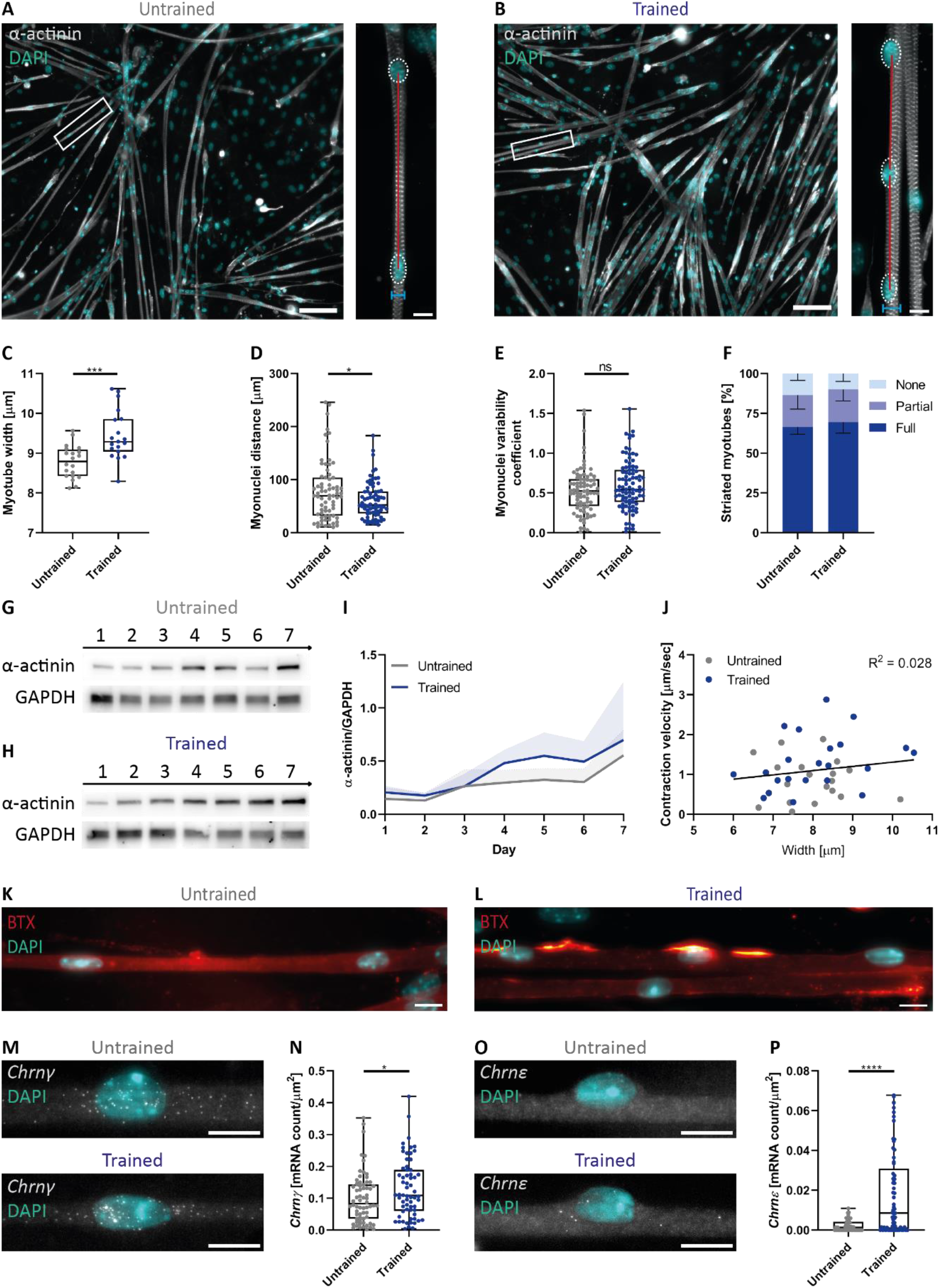
OptoTraining facilitates morphological maturation. **A, B)** Representative fluorescent images to assess myotube morphology of untrained and trained cultures (grey: α-actinin; cyan: myonuclei; Scale bars: 100 μm). Magnifications (white boxes; Scale bars: 10 μm) illustrate quantified tissue quality parameters. Myotube width (blue lines) and myonuclei spacing, density and uniformity (red lines). **C, D, E)** Boxplots showing **C)** myotube width, **D)** myonuclei distance and **E)** myonuclei variability coefficient at day 4 of differentiation. **F)** Percentage of striated myotubes (dark blue: fully striated; blue: partially striated; light blue: non-striated). **G, H)** Western blots of α-actinin and GAPDH (loading control) protein expression over 7 days of myotube differentiation. **I)** Graph showing temporal α-actinin expression normalized to GAPDH. **J)** Contraction velocity plotted relative to cell width of trained and untrained myotubes. R2 (0,028) was computed for the whole data set of untrained and trained myotubes via two-tailed Pearson’s correlation. **K, L)** Fluorescence images of α-bungarotoxin (BTX, red) showing acetylcholine receptor clusters and myonuclei (DAPI, blue). **M-P)** Chrnγ and Chrnε smFISH probes (grey) and DAPI staining (cyan) in trained and untrained myotubes with plot showing Chrnγ and Chrnε mRNA count per μm2 myotube area. Scale bars: 10 μm.

As OptoTraining promotes morphological maturation, we aimed to further validate this finding by looking at acetylcholine receptors (AChR), neurotransmitter-gated ion channels at the neuromuscular junction. During myogenesis, AChR segments increase in size and change their composition from embryonic (γ) to adult (ε) subunits (Jin et al., 2008; Cetin et al., 2020). To detect AChR in our cultures, we first stained myotubes with α-bungarotoxin (BTX) at day 4 of differentiation. We observed an increase in AChR clusters around myonuclei from trained myotubes (**figure 3K, L**). To explore further this enhanced AChR clustering, we evaluated the expression of both gamma (*Chrnγ*-fetal) and epsilon (*Chrnε*-adult) subunits at the transcriptional level. Using single molecule RNA fluorescence in situ hybridization (smFISH; Pinheiro et al., 2021) at day 4 of differentiation, we observed a higher expression of *Chrnγ* and *Chrnε* for trained myotubes (**figure 3M-P**). The detection of adult AChR mRNAs further confirms enhanced maturation of trained myotubes (Bakooshli et al., 2019) but most likely does not account for the accelerated contraction dynamics considering contractions are optogenetically driven.

### Increased contractile demand triggers an upregulation of neo-Myh8 and fast Myh

We hypothesized that the faster contractile dynamics after long-term training could be due to Myh switching. The pattern of Myh expression during development follows a sequential transition (Agbulut et al., 2003; Brown et al., 2012) from developmental (emb-Myh3 and neo-Myh8) to adult Myh isoforms (slow-Myh7 or fast Myh (fMyh: fast-Myh2, fast-Myh1, fast-Myh4; **figure 4A**)). We first investigated the expression of the developmental emb-*Myh3* and neo-*Myh8* genes at the single cell level. Due to the sequence homology of Myh genes (Weiss et al., 1999), we designed smFISH RNA probes against pre-mRNAs to ensure detection specificity. Whereas the RNA levels of emb-*Myh3* do not change after long-term exercise (**figure 4B, C**), we observed a significant upregulation of neo-*Myh8* (**figure 4D, E**) and downregulation of fast-*Myh7* (**figure 4F, G**) in trained myotubes.

**Figure 4).**
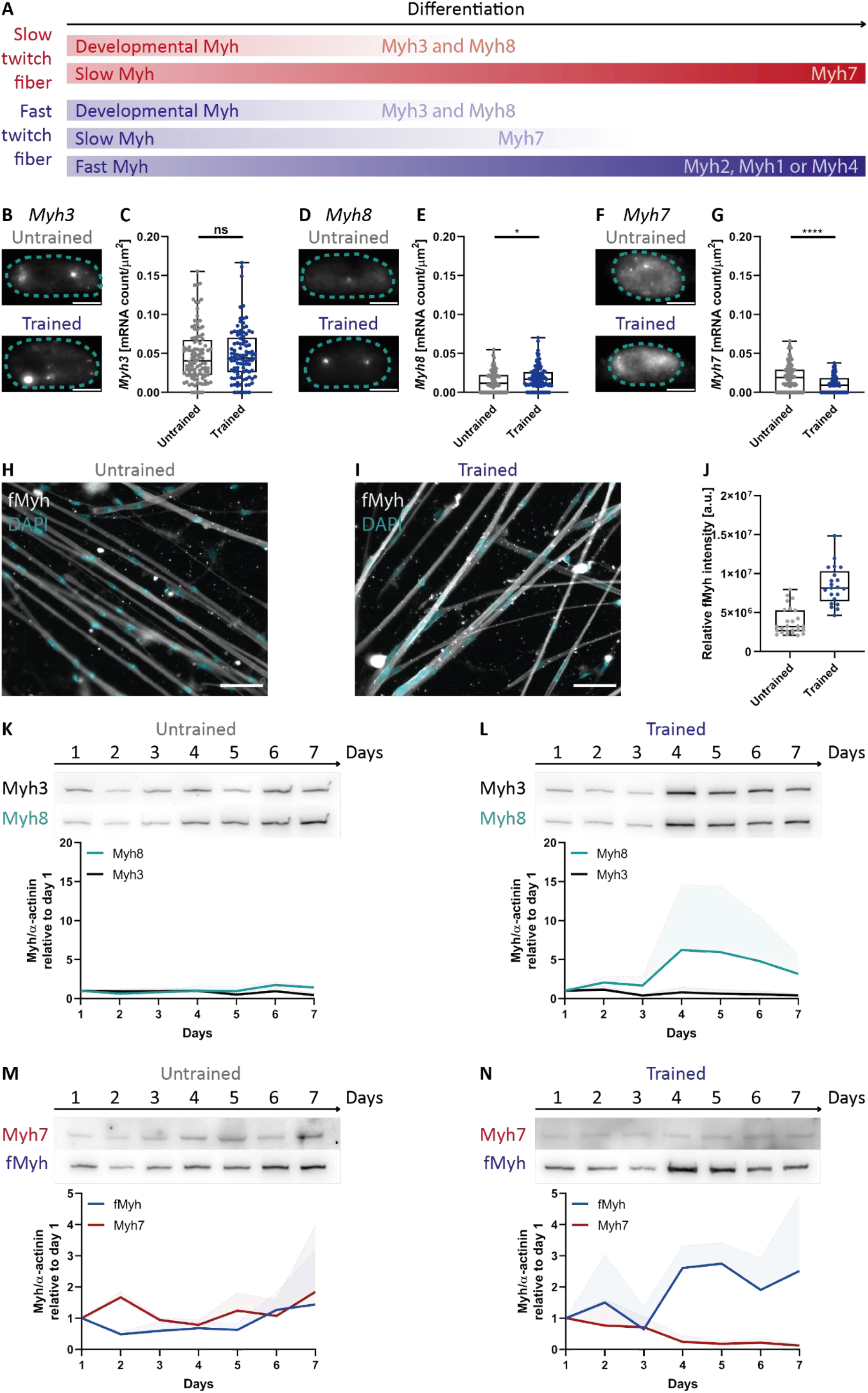
Long-term mechanical training upregulates expression of neo-Myh8 and fMyh isoforms. **A)** Simplified scheme showing temporal expression patterns of Myh isoforms for slow and fast twitch fiber types. **B-G)** Left: smFISH for Myh3, Myh8 and Myh7 pre-mRNA expressed in myonuclei (cyan dotted outline) of untrained and trained myotubes. Right: boxplots of RNA expression per myonucleus area. **H, I)** Immunofluorescence images of untrained and trained myotubes stained for fMyh. **J)** Measured fluorescence intensity of fMyh per myotube. **K-N)** Western blots of developmental and adult Myh isoforms (top) with associated graphs showing relative expression profile of respective Myh isoforms normalized to α-actinin (bottom), which was used as a muscle-specific loading control. Scale bars B and D: 5 μm; Scale bars F and G: 50 μm.

As Myh8 upregulation is characteristic of fast fiber type specialization (Schiaffino et al., 2015), we stained myotubes for fMyh (**figure 4F, G**). We measured a higher fluorescent intensity for trained cultures (**figure 4H**), suggesting a shift towards a fast fiber phenotype. To confirm this slow-to-fast fiber type switch, we investigated the temporal expression pattern of developmental and adult Myh isoforms over 7 days using western blots. Protein expression levels were normalized to α-actinin to account for sarcomerogenesis. Over the 7 days of differentiation, untrained cultures express both developmental and adult Myh isoforms with only minor fluctuations in relative expression (**Figure 4J, K**). In contrast, trained myotube cultures alter their Myh expression profile. For developmental Myh isoforms, emb-Myh3 decreases slightly whereas neo-Myh8 increases after the onset of training (Day2; **figure 4K**). For adult Myh isoforms, levels of slow-Myh7 decrease and are accompanied by a rapid upregulation of fMyhs (**figure 4L**), mainly fast-Myh1 (**supplementary figure 4A, B**). The predominant expression of neo-Myh8 and fMyh in trained cultures explains the increased contractile velocity and is characteristic of fast twitch fiber establishment during development (Johnson et al., 2019).

## Discussion

Fiber type composition confers muscle the appropriate kinetics to serve their contractile demands. This is particularly relevant during development (Agbulut et al., 2003) or exercise (Qaisar et al., 2016), in which fiber types can switch. In this study, we provide an experimental approach to accelerate myotube maturation and promote fiber type switching in a dish through *in vitro* exercise. Using an optogenetic setup compatible with cell culture, we induce myotubes to contract according to specified training protocols over several days. Long-term exercise of myotubes results in faster contraction dynamics and resistance to fatigue. By monitoring RNA and protein levels of developmental and adult Myh isoforms, we observed the premise of fiber type switching in trained myotubes with the upregulation of neo-Myh8 and fMyh. The accelerated contraction dynamics are also accompanied by faster muscle cell maturation at both structural and transcriptional level. Overall, we provide an *in vitro* strategy to model fiber type switching during development, exercise, or disease.

Submitting myotubes to mechanical stretch is a widely used approach to induce muscle contractions. Various molecular and biochemical alterations, including improved maturation, increased myotube size, and alignment, have been reported (Ren et al., 2021). Optogenetic stimulation of C2C12, which under standard culture conditions do not contract, facilitates maturation of striated, contractile myotubes (Asano et al., 2015). Using pharmacological treatments, the development of contractility has been linked to an increase of intracellular calcium levels that promote sarcomere assembly (Sebille et al., 2017; Chapotte-Baldacci et al., 2022). To further assess developmental maturity and define fiber type, Khodabukus et al. have characterized twitch dynamics of electrically stimulated 3D engineered myobundles (Khodabukus et al.; 2019). Trained muscle cells possess faster relaxation kinetics accompanied by higher force production, decreased fatigability, and increased glucose metabolism. However, no changes in glucose and mitochondria protein levels were detected, suggesting only a partial shift towards a fast fiber type. Most studies use intermittent stimulation patterns to mimic neuronal input-like stimuli. A complete fast fiber type shift may require a faster stimulation frequency, found *in vivo* (Hennig & Lømo 1985), and longer culture times.

Improvements of *in vitro* muscle cultures would provide greater control over fiber type switching. Aligning and attaching muscle cells is a strategy to prolong and mature the cultures allowing to recapitulate the full transition from fetal to adult Myhs. Other cell types could also be co-cultured to influence the expression of certain fiber type and the coordinated expression of Myh within a single myofiber. Myotubes cultured without other cell types adopt a “default path” towards a slow fiber type. However, firing frequency of motor neurons during development determines myofiber type (Hennig & Lømo 1985), and denervation leads to uncoordinated Myh expression (Dos Santos et al., 2020). Both processes could be recapitulated in optogenetically-stimulated motor neurons co-cultured with muscle. Such systems would be particularly relevant to investigate myonuclei coordination and the mechanisms of fiber type switching at several cellular levels (genetic, structural, functional, and metabolic).

Controlling fiber type in cultured muscle systems will also be relevant for disease modelling when using patient-derived induced pluripotent stem cells (iPSCs). As many myopathies preferentially affect certain fiber types (Talbot & Maves, 2016), biological investigation and drug testing should be performed accounting for fiber type composition. It will primarily be interesting to observe if the susceptibility of certain fiber type to muscle diseases is conserved *in vitro*. This would provide drug development models for therapies aimed at promoting fiber switching to the less afflicted fiber type. For example, compounds found to preserve muscle integrity in Duchenne Muscular Dystrophy (DMD) promote the less affect slow muscle phenotypes. Additionally, exercise protocols can be personalized to prevent or delay fiber switching in muscle disorders worsened by fiber composition alterations.

## Material and Methods

### Primary mouse myotubes *in vitro* culture

All procedures using animals were approved by the institutional ethics committee and followed the guidelines of the Portuguese National Research Council Guide for the care and use of laboratory animals.

*In vitro* myotubes were generated as described previously from primary mice myoblasts (Pimentel et al., 2017). Shortly, myoblasts were isolated from C57BL/6J mice 5-7days after birth. The tissue was minced and digested using 0.5mg/ml collagenase (Sigma) and 3.5 mg/ml dispase (Roche) in Phosphate-buffered saline (PBS). The cell suspension was then filtered and cells were plated in Iscove’s Modified Dulbecco’s Medium (IMDM, Thermo Scientific, 37°C, 5% CO2). After 4 hours, the medium was collected containing non-adherent myoblasts. Cells were then centrifuged (1400 rpm, 5 min) and resuspended in growth medium (IMDM + 20% Fetal Bovine Serum (FBS) + 1% chick Embryo Extract + 1% Penicillin/Streptomycin). Cells were plated onto IBIDI dishes coated with 1:100 matrigel (Corning Inc.) at a density of 200000 cells/ml. Differentiation was initiated after 4 days by changing the medium to differentiation medium (IMDM + 10% Horse serum (HS) + 1% Penicillin/Streptomycin). At day 1 of differentiation, a thick layer of 150 ul matrigel (1:1) was added on top of forming myotubes and the medium was supplemented with 100ng/ml agrin. Cells were cultured up to 7 days at 37°C and 5% CO2.

### OptoTraining

At day 0 of differentiation, cells were infected with pACAGW-ChR2-Venus-AAV9 (Addgene). 0.5 ul were added to 500 ul differentiation medium. Cells were incubated O/N at 37°C and 5% CO2. Cultures were washed once with PBS, before matrigel and differentiation medium were added as described before. At day 2, OptoTraining was initiated using the OptoPlate. Trained cell were submitted to specific temporal light pattern (50 ms light pulses at 2, 5, or 10 Hz) for 8 hours per day. ChR2 expressing control cells were kept in the dark.

### Cell fixation and immunostainings

Cells were washed 1 x with PBS and fixed for 10 minutes with 200 μl of 4% PFA. Subsequently, dishes were washed 2 x with PBS. If needed, cells were incubated with 1:50 Alexa Fluor 488 α-bungarotoxin (BTX, Invitrogen) in PBS for 20 min at RT. Cells were permeabilized with 0.5% tritonX100 in PBS for 5 minutes and blocked with 10% Goat serum in PBS with 5% BSA (blocking buffer, BB) for 1 hour. Primary antibodies were diluted in 200 μl BB with 0.1% saponine and incubated overnight at 4°C. Dishes were washed with PBS 3 x for 5 minutes under agitation. Secondary antibodies (1:400) together with DAPI (1:10000) were incubated in BB + 0.1% saponine for 1 hour and subsequently washed 3 x for 5 minutes with PBS under agitation. 150 μl of fluoromount-G (SouthernBiotech) were added per dish and dried for 24 hours at 4°C. All use antibodies are listed in table 1. Images were acquired using an inverted widefield fluorescence microscope (Zeiss Cell Observer).

**Table 1).** List of primary antibodies

| Target | Dilution immunostaining | Dilution western blots | Company | Catalog number |
| --- | --- | --- | --- | --- |
| Alpha actinin | 1:200 | 1:1000 | Sigma Aldrich | A7732 |
| fMyh | 1:200 | 1:5000 | Santa Cruz Biotechnology | sc-32732 |
| Myh1 | - | 1:1000 | Thermo Fisher Scientific | # PA5-117077 |
| Myh2 | - | 1:5000 | Santa Cruz Biotechnology | sc-53095 |
| Myh3 | - | 1:5000 | Developmental Studies Hybridoma Bank | F1.652 |
| Myh7 | - | 1:250 | Developmental Studies Hybridoma Bank | A4.951 |
| Myh8 | - | 1:10000 | Thermo Fisher Scientific | PA5-72846 |
| Filamin C | 1:200 | 1:5000 | MyBioSource | MBS2026155 |

### Structural analysis of cellular maturation

Stained images were pre-processed using FIJI (Schindelin et al., 2012) to apply z-projection, subtract background and binarize images stained with DAPI and α-actinin. The proportional area of myotubes in each image was calculated using an in-house MATLAB (R2019a; MathWorks, Natick, MA) code to determine the proportion of regions with α-actinin fluorescence in binarized images. The total area of myotubes was then divided by the total length of the centerline skeletons of the myotube regions to estimate the average myotube width for each image. DAPI stained nuclei residing within regions of α-actinin fluorescence were designated as myonuclei. The sum of myonuclei in each image was determined using MATLAB code and normalized to calculate myonuclei per mm^2^.

Myonuclei distribution was measured in MATLAB by selecting myotubes at random and manually labelling nuclei. This spatial distribution data was then used to calculate the mean distance between neighboring nuclei in each myotube. Uniformity of myonuclei distribution was determined using the *coefficient of variation* (standard deviation in distance divided by mean distance between nuclei for each recorded myotube). The percentage of randomly selected myotubes with full, partial (<50%) and no striations were recorded manually.

FIJI macros for pre-processing images and MATLAB code for segmentation and calculation of metrics is available on Github (https://github.com/dhardma2/MyoChip/Opto).

### Western blots

Dishes with myotubes were washed 3 x with cold PBS on ice. Cells were removed from ice and 200 ul of lysate buffer (1% SDS in 100mM TRIS-HCl, pH 8) added to dish. Cells were scrapped and transferred into 1.5 ml Eppendorf. To remove matrigel, cells were spun down for 5 minutes at 14000 rpm. The supernatant was collected and stored at -80°C. For western blots, cell lysate protein concentrations were measured using the BCA kit (Pierce). Samples were mixed with 1:4 lammeli buffer and boiled at 95°C for 5 minutes. The same amount of sample (10 or 30 ng/ml) were loaded onto 4-15% pre-cast Bis-tris gel (Invitrogen) and run at 100V. Proteins were transferred onto nitrocellulose membrane for 75 minutes at 100V. Subsequently, membranes were blocked for 1 hour in blocking buffer (BB) using 5% non-fast dry milk in TBS-T (TRIS-buffered saline with 0.1% Tween 20). Primary antibodies in BB were incubated O/N at 4°C. Membranes were washed 3 x with TBS-T under agitation and then incubated with HRP-conjugated secondary antibodies for 1 hour at room temperature. Membranes were washed 3 x in TBS-T under agitation, visualized using ECL reagent (Peirce) and imaged using Amersham ImageQuant 800 Western blot imaging system. Quantification was done in Image Lab. All used antibodies are listed in Table 1.

### Single molecule FISH

For *Chrnγ, Chrnε*, and *filamin C*, mRNA probes were designed to align with the coding sequence of the mRNA of interest using the Stellaris probe designer (sequences listed in Table 2). For myh isoforms, smFISH probes were designed for pre-mRNA sequences containing introns and exons to ensure high specificity. All probes were coupled to Quasar570. The dried oligonucleotide probe was dissolved in 400 ul RNase free water (Invitrogen) to a stock concentration of 12.5 μM. Culture dishes with myoblasts were washed with RNase-free PBS (Ambion) and fixed 10 minutes at RT in fixation buffer (10% formaldehyde solution, Sigma Aldrich, in nuclease free water). Cells were washed 2 x and permeabilized O/N in 70% ethanol at 4°C. For hybridization, dishes were washed with wash buffer (1x saline-sodium citrate, SSC, Sigma-Aldrich, in 10% deionized formamide, Ambion) for 5 minutes. 1.25 μM smFISH RNA probe in 10% formamide, 1% dextran sulfate (Sigma Aldrich) in 2x SSC and incubated O/N at 37°C. Myotubes were washed 2 x in wash buffer (containing 50 ng/ml DAPI at second wash) for 30 minutes at 37°C. 1 ml 2x SSC was incubated for 5 minutes at room temperature. Cells were then mounted using 150 μl Vectashield Antifade Mounting Medium (Vector Laboratories) and stored at 4 °C. Cells were imaged within one week using an inverted widefield fluorescence microscope (Zeiss Cell Observer). All images were processed in Fiji. Z-stacks of myofibers were projected to maximum intensity, background was subtracted and images were binarized. For *Chrnγ, Chrnε* and *filamin C*, mRNA signals were counted within the perinuclear region (myonuclei ±50 μm along the myotube). For myh isoforms, the area per myonuclei was computed.

**Table 2).** List of FISH probes

### Contractility assay and PIV analysis

To determine functional properties of untrained and trained cells, we measured myotube contraction velocity and fatigability at day 4 of maturation. To do so, we performed time-lapse imaging with high temporal resolution (20ms/frame) using a Zeiss Cell Observer SD microscope (63x oil immersion objective Plan-Apochromat 63x, NA M27). Single striated myotubes with nuclei at the periphery were chosen. First, cells were imaged without photostimulation. Subsequently, cells were submitted to continuous blue light illumination (470nm). We computed contractile parameters over time periods of 2 seconds in MATLAB using PIVlab, an image-based PIV analysis software (Thielicke et al., 2014). Due to heterogeneity of contraction along the muscle cell, myonuclei were used as reference points to measure displacement and instantaneous velocity per myotube. The program divides images into small interrogation windows (FFT window deformation, interrogation area phase 1: 64 pixel; phase 2: 32 pixel). Via maximum correlation method the local displacement of two consecutive windows is computed. From this, the program calculates quantitative parameters (e.g. velocity, displacement) of myotube movement and displays vector and velocity magnitude fields. To assess fatigability, myotube contractions were induced via continuous light stimulation. And the time was measured until cells stopped moving.

### Displacement curve analysis

A Fast Fourier Transform (FFT) was applied to myonuclei displacement curves in MATLAB to determine the frequency of a representative sine wave for the motion of each cell after stimulation. To assess variation in muscle cell twitch motion over time, in-house MATLAB code was used to find the maxima and minima of single cell displacement curves recorded over 2 seconds. The amplitude for each twitch was calculated and its contraction and relaxation time were computed. The code used for PIV data analysis is available on Github (https://github.com/dhardma2/MyoChip/Opto).

### Statistical Analysis

Statistical analysis was carried out in GraphPad Prism (using unpaired parametric t-test). The distribution of data points is expressed as mean ± SD for three or more independent experiments. Outliers were identified using ROUT method (Q=1).

## Acknowledgements

We thank all partners of the MyoChip consortium and all members of the Edgar Gomes and the Claudio Franco laboratory. We thank Afonso Malheiro and Judite Costa for support and discussions. We thank António Temudo, Ana Nascimento, Aida Lima and José Rino, from the Bioimaging facility at iMM, for imaging assistance.

## Author contribution

K.H carried out experiments and analyzed data. D.H performed the computational analysis. D.B. designed and produced the Optoplate. I.M. isolated primary myoblasts. K.H. and W.R conceived and designed the experiments. K.H, M.O.B, E.R.G. and W.R. wrote the manuscript with assistance from other authors and all authors participated in the critical review of the manuscript.

