## Supplementary figures for "Generating fast-twitch myotubes *in vitro* using an optogenetic-based, quantitative contractility assay"

### Supplementary information

### Supplementary figures

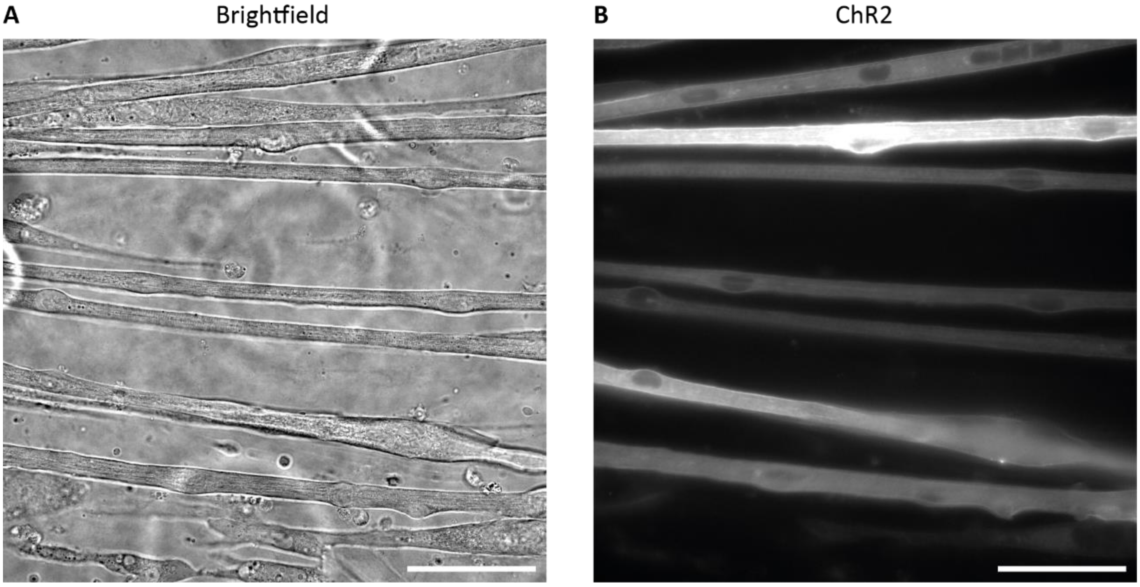


***Supplementary figure 1) AAV9-pACAGW-ChR2-Venus-AAV infection leads to high percentage of myotubes expressing ChR2. A)*** *Brightfield image of myotubes at day 4 of differentiation.* ***B)*** *Fluorescent image of ChR2. Scale bar: 50 µm.*

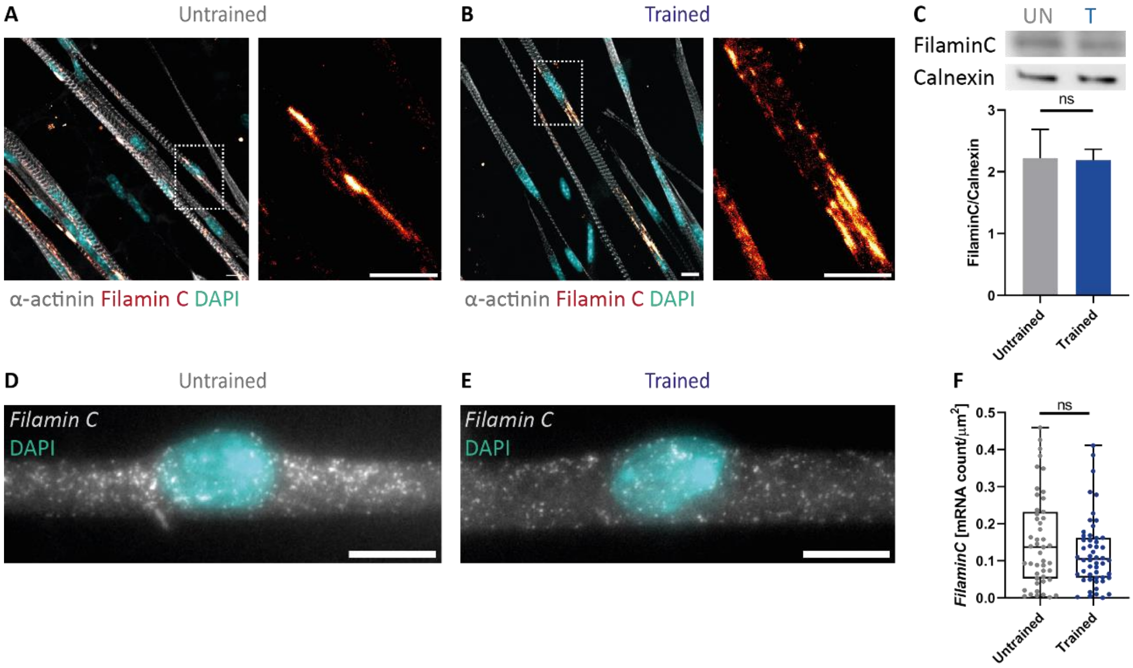


***Supplementary figure 2) OptoTraining does not induce sarcomeric damage. A, B)*** *Immunofluorescent images of filamin C showing sarcomeric scaring (hot red) in untrained and trained myotubes at day 4 of differentiation (cyan: DAPI; grey: α-actinin).* ***C)*** *Western blots and quantification of filamin C and calnexin (loading control) protein expression.* ***D, E)*** *smFISH probes for filamin C expressed in myotubes.* ***F)*** *smFISH quantification of filaminC mRNA counts. Scale bars: 10 µm.*

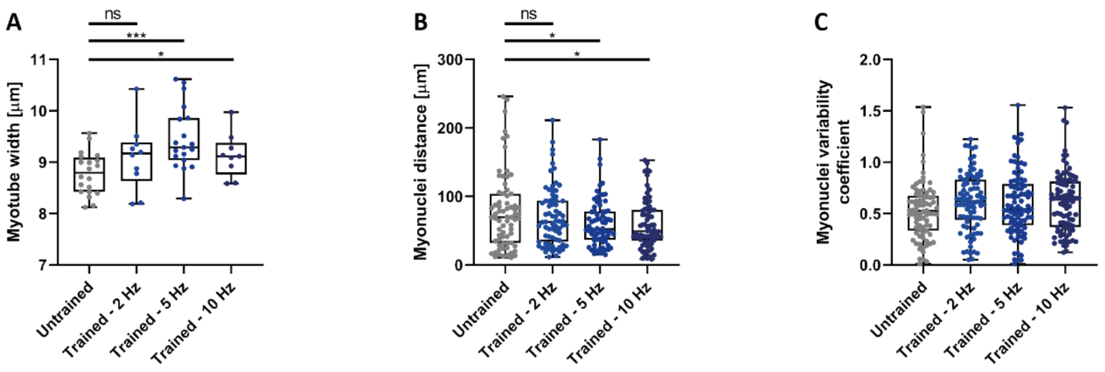


***Supplementary figure 3) Myotubes adapt morphology to distinct stimulation frequencies. A, B, C)*** *Box-plots showing **A)** myotube width, **B)** myonuclei spacing distances and **C)** uniformity coefficient at day 4 of differentiation for untrained myotubes and cultures stimulated at 2, 5 or 10 Hz.*

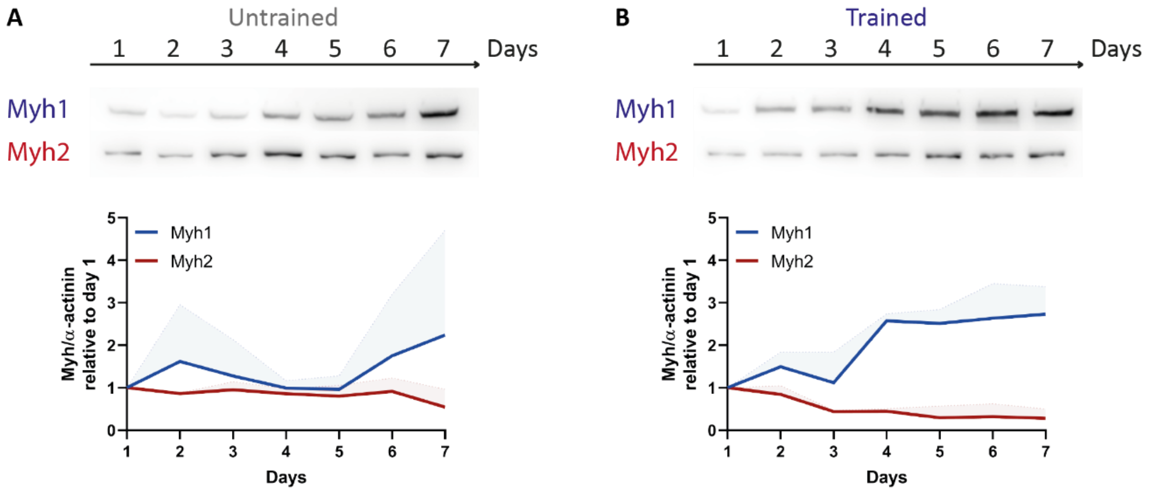


***Supplementary figure 4) Upregulation of fMyh in trained muscle cultures is due to increase in fast-Myh1 protein expression. A, B)*** *Western blots of fast-Myh1 and fast-Myh2 protein expression over 7 days in untrained and trained muscle cultures. Graphs show temporal expression profile of Myh isoforms relative to α-actinin, which was used as a loading control to account for sarcomerogenesis, and normalized to day 1 of differentiation.*
